# Extraction-free colorimetric RT-LAMP for SARS-CoV-2 RNA detection from saliva

**DOI:** 10.1101/2024.08.09.607279

**Authors:** Yuying Yang, Xing Liu, Songyan Tie, Xin Li, Xin Yang, Jianzhong Cao

**Author notes:** Corresponding author: Prof. Jianzhong Cao, College of Chinese Medicine, Hunan University of Chinese Medicine. 300 Xueshi Road, Changsha, Hunan Province, P.R. China. Authors email: Yuying Yang, Xing Liu, Songyan Tie, Xin Li, Xin Yang.

## Abstract

**Aim:** To develop a rapid and easy diagnostic assay for detection of SARS-CoV-2 RNA presented in saliva samples.

**Method:** The color-based RT-LAMP was used to detect nucleocapsid (N) gene of SARS-CoV-2 without RNA extraction from saliva.

**Results:** RNA spiked saliva can be used directly for cDNA synthesis after heat inactivation of the saliva and diluted 2-fold with either water or PBS. For both PCR and LAMP, 20% of saliva did not have obvious effect on the reaction. Saliva did not interfere with RT-LAMP when the volume of saliva was less than 20% of the total volume. The sensitivity of the RT-LAMP reached to 1.034×10^-5^ ng/µl (475 copies/μl). The RT-LAMP assay was validated by testing 20 RNA spiked saliva samples. The assay specificity was similar to that of data without saliva.

**Conclusions:** The RT-LAMP colorimetric assay can be used as a screening method with the advantages of being rapid, easy to use than the qRT-PCR.

## Introduction

The quantitative real-time polymerase chain reaction (qRT-PCR) has become the gold standard for diagnosis of severe acute respiratory syndrome coronavirus 2 (SARS-CoV-2) infection since the outbreak of COVID-19 pandemic(WHO, 2020). The current qRT-PCR assay detects RNA of either nucleocapsid (N) gene or N and ORF1ab gene(WHO, 2020). The lung or nasopharyngeal swabs have been used as the major sources of sampling and the collection of swabs requires individuals going to indicated hospitals or stations, increasing risks of sampling-associated infection. In addition, the swabs cannot be used directly for PCR, but require RNA purification and cDNA synthesis before PCR, resulting in hours to days to get the result and thence the infected people might miss the optimal treatment window. Besides, the PCR assay requires expensive PCR machine, reagents, and specialized technicians, so increases the cost of the diagnosis. Therefore, family based, low cost, and easy to use assays are in demand worldwide not only for SARA-CoV-2 but other viral infections(Byrne et al., 2020).

As an alternative to swab, saliva has been demonstrated to be a reliable source of SARS-CoV-2 RNA for diagnosis(Altawalah, AlHuraish, Alkandari, & Ezzikouri, 2020; Azzi et al., 2020; Byrne et al., 2020). Previous studies revealed that viral RNA in swabs showed a dynamic change during the day, while viral RNA in saliva remained relatively stable during the same period, indicating saliva was a more reliable source of RNA than swabs(Chatterjee et al., 2020; To et al., 2020). Thence, saliva based qRT-PCR has been developed for diagnosis of SARS-CoV-2 infection. However, all PCR based methods have similar issues in terms of cost and time. Contrary to PCR, loop-mediated isothermal amplification (LAMP) is a simple, low cost, easy to perform nucleic acid detection method(Chow, Chan, Tam, Zhao, Yao, Fung, Cheng, & Lo, 2020; Gandelman, Jackson, Kiddle, & Tisi, 2011). Particularly, the result of LAMP assay can be obtained by observing the color change of the reaction solution, avoiding expensive equipment and tedious calculation.

Previous reports using LAMP for COVID-19 diagnosis used purified RNA from saliva or swabs and achieved better results both in sensitivity and reliability(Chow, Chan, Tam, Zhao, Yao, Fung, Cheng, & Lo, 2020; Dao Thi et al., 2020; Ooi et al., 2022). However, most of the reports used primers targeting N and ORF1ab gene, thus single-positive results were seen when RNA concentration was low(Chow, Chan, Tam, Zhao, Yao, Fung, Cheng, Lo, et al., 2020; Dao Thi et al., 2020; Zhang et al., 2020). We reasoned that saliva contains not only the virion particles, but also the cell debris or vesicles released from infected cells. Thus, the viral subgenomic RNA would contribute a lot to the total viral RNA. As N gene presented in every subgenomic RNA and genomic RNA, it also is the less mutated gene of SARS-CoV-2, the use of N gene as detection target would be a more reliable way for family-based diagnosis.

In this study, we developed a rapid, ready-to-use, colorimetric RT-LAMP assay that can be applied within a hospital setting or a family for detection of SARS-CoV-2 RNA using inactivated saliva sample.

## Materials & Methods

### 1. Reagents

The plasmid containing N gene of SARS-CoV-2 was kindly provided by Professor Xinyi Ge, College of Life Science, Hunan University, Changsha, China. The plasmid pCE, the plasmid mini-isolation kit, the cDNA synthesis kit, and T7 High Yield RNA Transcription kit were purchased from Vazyme (Vazyme Biotech Co., Ltd., Nanjing, China). The RNA purification kit was purchased from AG (Accurate Biotechnology Co., Ltd., Hunan, China). The chemical competent cell DH5alpha was purchased from Weidi (Shanghai Weidi Biotechnology Co., Ltd, Shanghai, China). The PCR amplification kit was ordered from AG (Accurate Biotechnology Co., Ltd., Hunan, China). The WarmStart Colorimetric LAMP 2X Master Mix kits were purchased from NEB (NEB, Beijing, China). Other molecular biology reagents were commercially available. The concentration of DNA or RNA was determined by NanoDrop (Thermo Fisher Scientific Co., Ltd., USA). The qPCR or fluorescent RT-LAMP was performed on LightCycler®480/96 (Roche Co., Ltd., Switzerland). The PCR primers for N gene were NF (GTTCACCGCTCTCACTCAAC) and NR (TTCGTCTGGTAGCTCTTCGG). The PCR conditions were following the instructions of the kits and were run for 35 cycles in regular PCR or 40 cycles in qPCR.

### 2. Saliva collection and inactivation

The human saliva samples were collected from healthy individuals on campus of Hunan University of Chinese medicine with the consent of the volunteers. Each saliva sample was collected in a 1.5 ml Eppendorf tube and capped. After cleaning the surface with 70% ethanol, the tubes were incubated for 10 min at 95°C to inactivate saliva. Then the tubes were kept at room temperature for 5 min. To reduce the viscosity of the saliva, 500 µl of saliva was mixed with 500 µl of DNase-free and RNase-free water to give a saliva stock. This saliva stock was easy to handle with routine pipet tips. The stock solution was stored at -20°C before use.

### 3. RNA and DNA template preparation

The N gene of SARS-CoV-2 was amplified by PCR using primers NF1 and NR1 and cloned into pCE plasmid downstream of T7 promoter to generate plasmid pCE-N. To transcribe N RNA from pCE-N, one microgram of plasmid was added to a 1.5 ml tube containing transcription buffer. The final reaction volume was 20 µl. The transcription reaction was performed at 37°C for 30 min and then the plasmid DNA was digested with DNase I for 5 min at 37°C. The RNA was purified by RNA purification kit and eluted with 50 µl of DEPC treated water. After quantification of RNA, the RNA sample was used for cDNA synthesis or stored at -80°C before use for other experiments. For cDNA synthesis, 100 ng of RNA was used in 20 µl of cDNA synthesis reaction using random primers and incubated at 45°C for 30 min. The cDNA was used directly for downstream amplification.

To prepare stock saliva sample containing different concentration of DNA or RNA, inactivated saliva was spiked with DNA or RNA stocks to give 99.5 ng/μl of DNA (2.17×10^9^ copies/µl) or 10340.8 ng/μl RNA (4.75×10^11^ copies/µl) and the total volume was brought to 500 µl with water. When in use, the stock solution was 10-fold diluted with inactivated saliva to give desired DNA/RNA concentrations. The samples without saliva were prepared with DNase-free and RNase-free water.

### 4. LAMP assay design

The primers for LAMP were designed using NEB LAMP tool (https://lamp.neb.com/#!/) and synthesized by Wuhan Shenggong Co. The primers for PCR and LAMP were listed in Table 1. A 10 X LAMP primer stock was prepared by mixing 16 μM each forward inner primers/backward inner primers (FIP/BIP primer), 4 μM each loop primer, and 2μM each F3/B3 primer. Depending on the experimental design, each LAMP reaction contained 12.5μl WarmStart Colorimetric LAMP 2X Master Mix, 2.5µl 10 X LAMP primer stock, 2-10 μl template sample, and sterilized water to give total volume of 25 μl. LAMP reaction was incubated at 65°C for 30 minutes on a metal incubator or PCR machine. The color images of LAMP reaction were captured by cell phone and part of the samples were analyzed by agarose gel electrophoresis and the images were captured by ChemiDOC XRS+ (BioRad. Co., Ltd., USA). When fluorescent RT-LAMP was performed, the LAMP reaction was the same as above except 0.5µl LAMP fluorescent dye was included in the reaction and the reaction was performed at 65°C on a LightCycler®480/96 machine. The fluorescence intensity was measured at 60 second intervals.

**Table 1.**
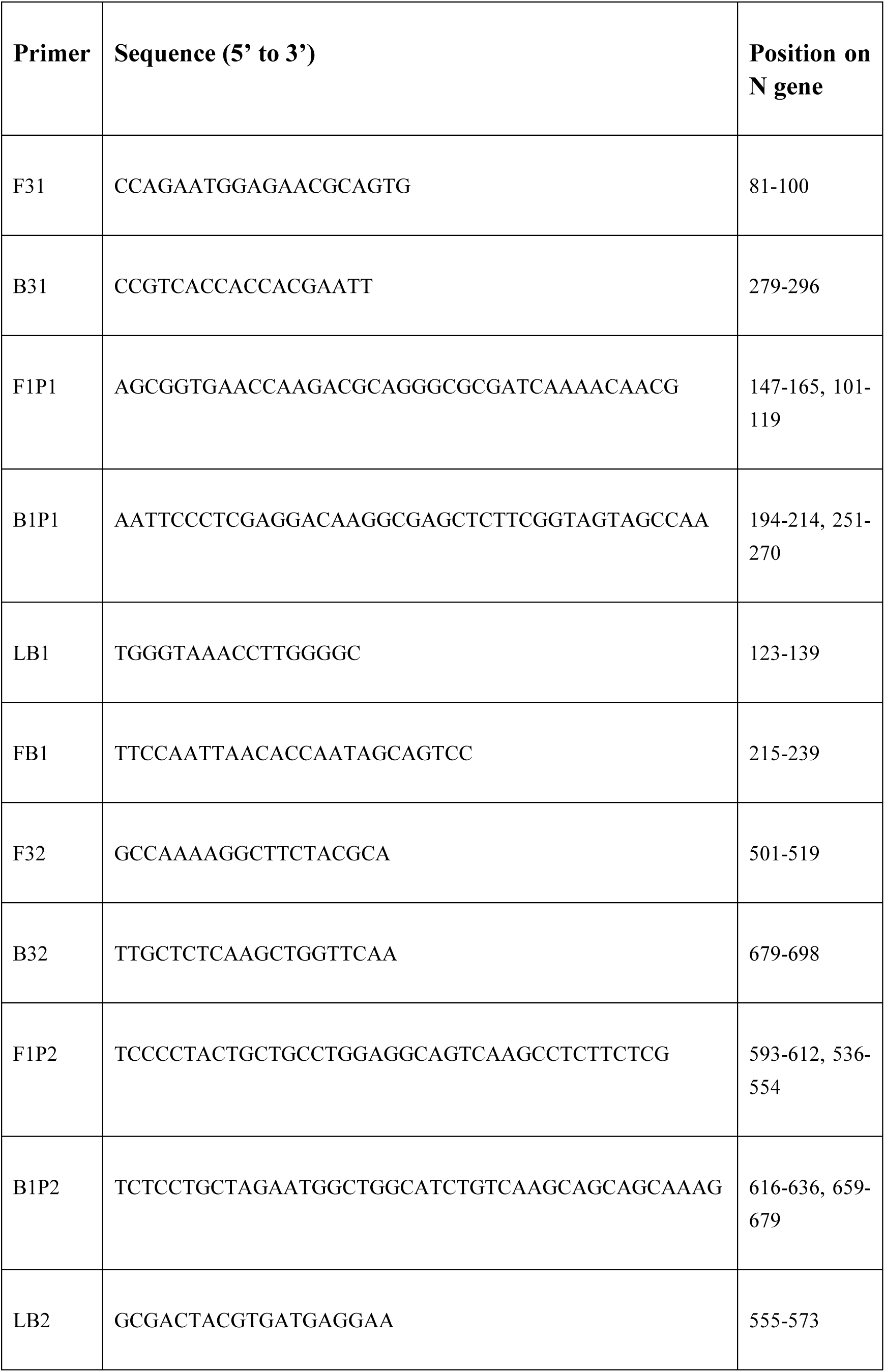
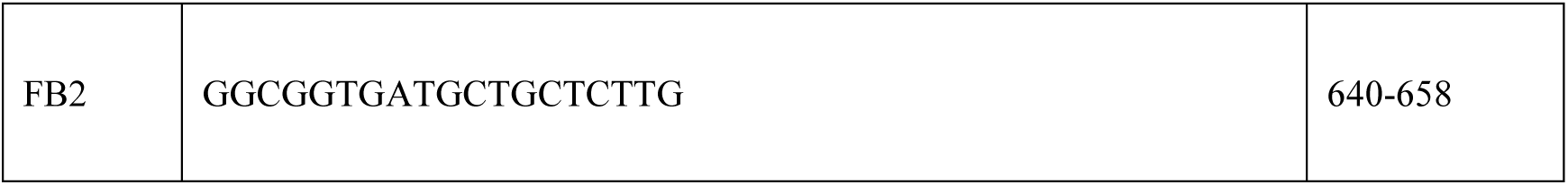
Primers for LAMP.

### 5. Statistical analysis

All data were expressed as mean ±SD (standard deviation).

## Results

### 1. Establishment of LAMP color assay

To develop a sensitive and reliable LAMP assay for COVID-19 saliva samples, we designed three sets of LAMP primers targeting the N gene of SARS-CoV-2 because N gene is the most abundant viral gene presented in the form of virial particles and viral infected cells which were released into the saliva as virions, cellular debris or vesicles. As shown in Table 1, the primer sets cover the different regions of the N gene, thus could provide similar sensitivity in the assay, avoiding the so called “single-positive” phenomena usually found in current RT-qPCR diagnostic assay. To test the primer specificity and sensitivity, we first cloned N gene into pCE vector and used it as template to perform routine LAMP assay. As shown in Fig. 1, all sets of LAMP primers effectively produced expected products after 30 min of incubation at 65°C and the color of the reaction changed from pink to yellow (Fig. 1c). The negative control showed no amplification and thence no change in color. The detection sensitivity reached to 1.034×10^-5^ ng/μl (475 copies/μl) with primer set #1 and #3 displayed the most sensitivity.

**Fig. 1.**
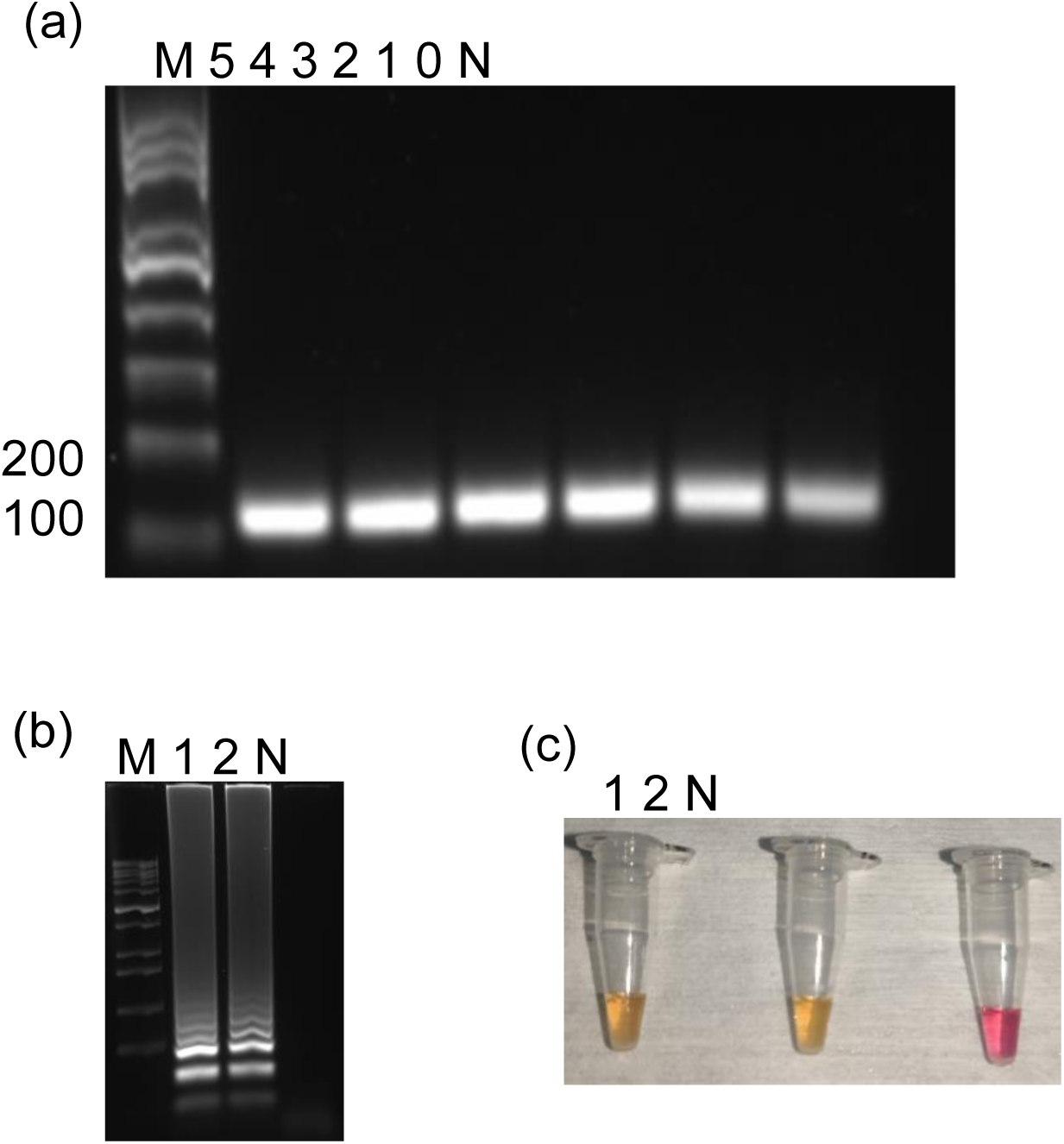
Development of PCR and LAMP assay. (a) SARS-CoV-2 N gene was amplified by PCR from plasmid pCE-N and analyzed by agarose gel. M, 100 bp DNA ladder. N, no DNA negative control. Number 0-5 indicated the amounts of DNA template were from 101 to 105 copies/ul. (b), Agarose gel analysis of LAMP products using two sets of LAMP primers, N was negative control. (c) Images of LAMP reaction as in (a).

### 2. Effect of saliva on cDNA synthesis

Because viral RNA has to be converted into cDNA for PCR or LAMP amplification, we first tested effect of human saliva on cDNA synthesis. To achieve this, saliva samples were collected from healthy human volunteers and inactivated at both 65 and 95°C for 30 or 5 min, respectively. Then, different volumes of the inactivated saliva was added to cDNA synthesis reaction. qPCR was used to amplify the cDNA to evaluate the effect of saliva on cDNA synthesis. As shown in Fig. 2, up to 8 µl of saliva (40% of final volume) did not affect the PCR amplification efficiency, indicating that saliva did not affect cDNA synthesis. Then, LAMP was performed using the cDNAs synthesized in the presence of saliva, and again, no obvious difference was found between cDNA samples synthesized in the presence or absence of saliva, indicating again inactivated saliva did not affect cDNA synthesis.

**Fig. 2.**
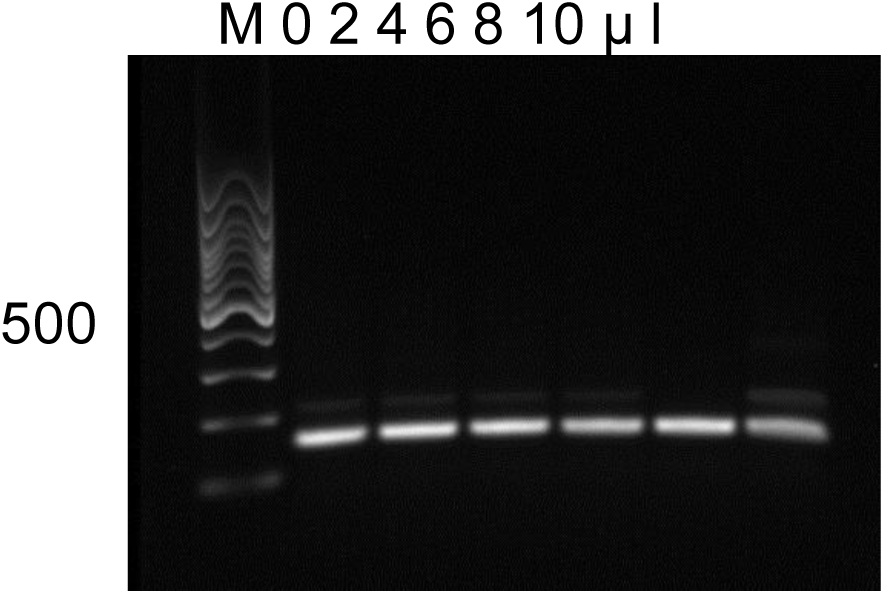
Effect of saliva on cDNA synthesis. RNA of N gene was reverse transcribed into cDNA in the presence of different amount of inactivated saliva. The final reaction volume was 20 ul and the volumes of the saliva were from 0 to 10 ul. After cDNA synthesis, 2 ul of cDNA was used for PCR in 25 ul of reaction and 5 ul of PCR products were ran on 1.2% agarose gel using TBE buffer. M, 100 bp DNA ladder.

### 3. Effect of saliva on PCR or LAMP

Though SARS-CoV-2 genome is RNA, both LAMP or PCR uses DNA template for amplification. To test whether saliva would interfere with LAMP or PCR, we performed LAMP and PCR in the presence of different volumes of inactivated saliva with fixed amount of N gene DNA. As shown in Fig. 3, saliva did not affect PCR (Fig. 3a, 3b) or LAMP (Fig. 3c, 3d) amplification when saliva was added up to 40% of final volume.

**Fig. 3.**
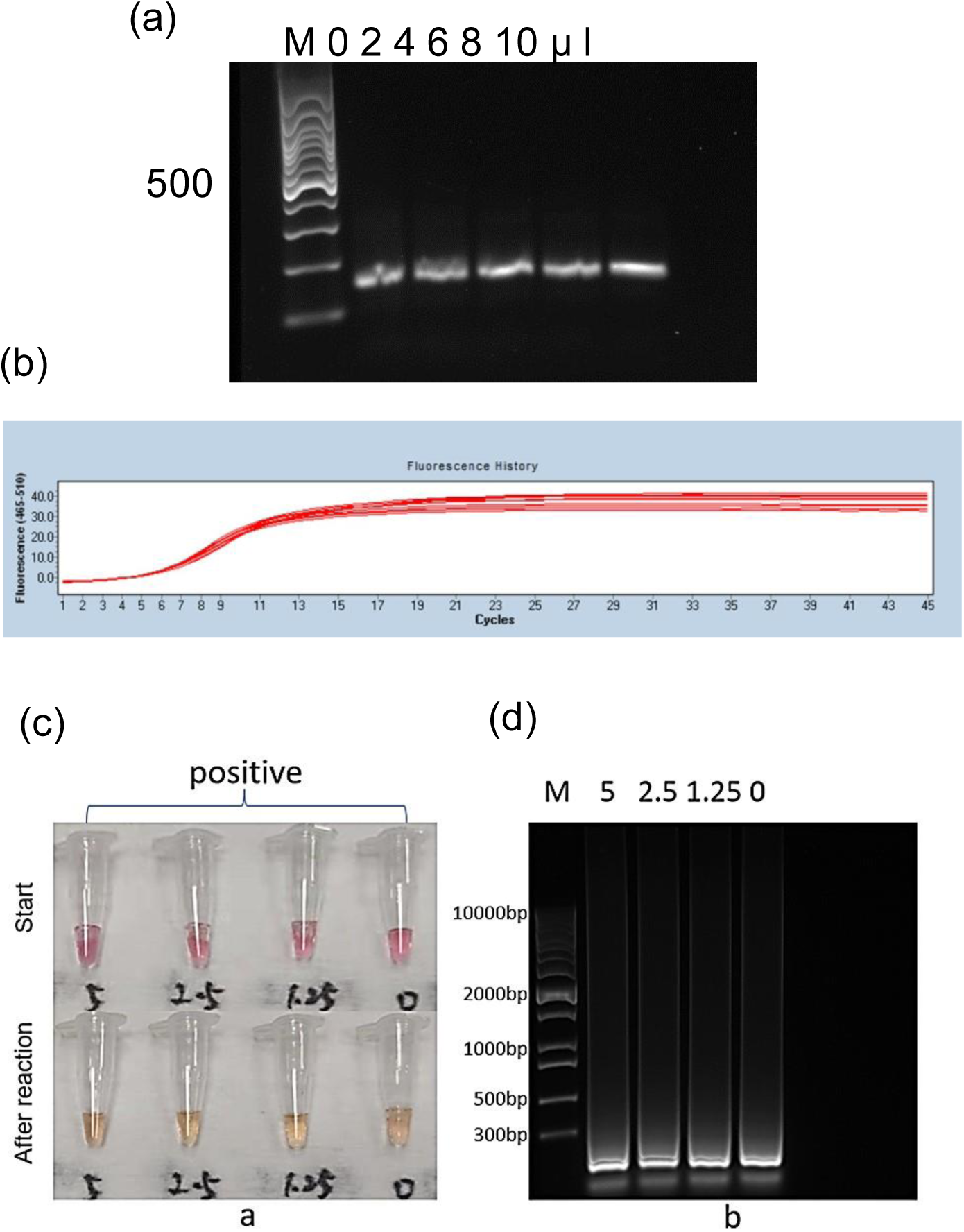
Effect of saliva on PCR or LAMP. PCR or LAMP was performed in the different amount of inactivated saliva. The plasmid pCE-N was used as DNA template. M, 100 bp DNA ladder. (a) Agarose gel analysis of PCR products. (b) PCR amplification. (c) LAMP result before and after reaction at 65oC for 30 min. (d) Agarose gel analysis of LAMP products.

### 4. Effect of saliva on RT-LAMP

Though above results did not find interference of saliva on cDNA synthesis, LAMP or PCR, we wondered whether saliva would interfere with RT-LAMP. To test this, we added different volumes of saliva to RT-LAMP reaction which contained 4.75×10^4^ copies of RNA and incubated the assay at 65°C for 30min. As shown in Fig. 4, compared with no saliva control, saliva did not affect RT-LAMP amplification when saliva volume was up to 40% final volume (Fig. 4a, 4b). Thus, these data clearly indicated that inactivated saliva did not interfere with either cDNA synthesis, PCR, LAMP, or RT-LAMP assay in our experimental settings.

**Fig. 4.**
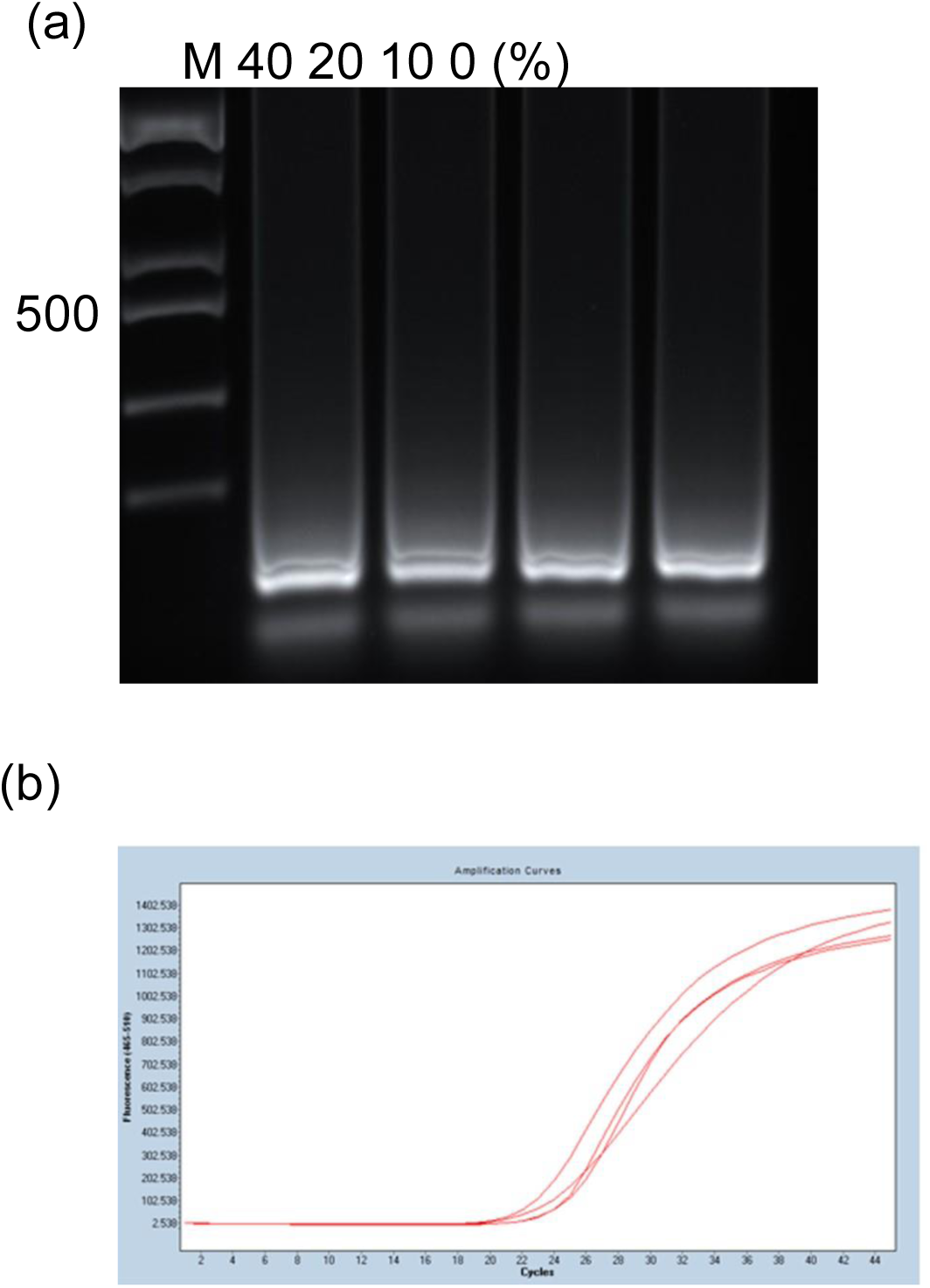
Effect of saliva on RT-LAMP. RT-LAMP was performed with RNA of N gene in the presence of inactivated saliva (0-40% final volume) at 65oC for 30min. M, DNA ladder. (a) Agarose gel analysis of RT-LAMP products. (b) Profile of RT-LAMP.

### 5. Saliva on the sensitivity of the RT-LAMP assay

To use RT-LAMP for detection of SARS-CoV-2 nucleic acid in saliva, the sensitivity is a concern. Thus, we performed RT-LAMP in the absence or presence of saliva that contained a serials of 10-fold diluted N gene RNA. As shown in Fig. 5, in the absence (Fig. 5a, 5b, 5c) or presence (Fig. 5d, 5e, 5f) of 40% saliva, the sensitivity of RT-LAMP reached to 4.75 X 10^3^ copies/reaction. So, 40% of inactivated saliva did not affect the sensitivity of the RT-LAMP.

**Fig. 5.**
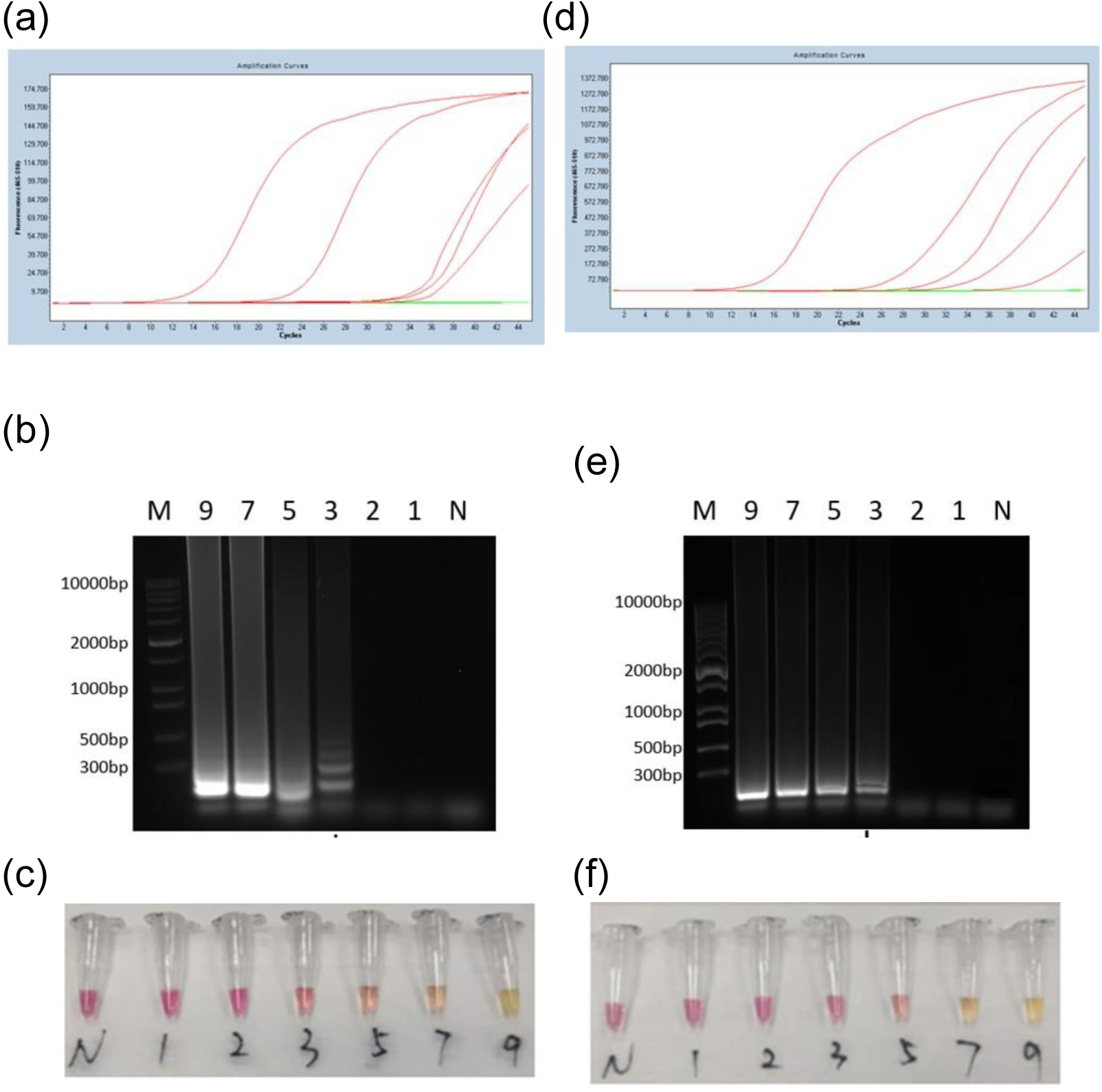
Effect of saliva on the sensitivity of the RT-LAMP. RT-LAMP was performed in the absence of saliva (a, b, c) or in the presence of 40% inactivated saliva (d, e, f) with different amount of N gene RNA at 65oC for min. (a, d) Profile of RT-LAMP in the absence of saliva. (b, e) Agarose gel analysis of RT-LAMP products. (c, f) Representative images of the RT-LAMP reactions. N, no RNA control; number 1, 2, 3, 5, 7, 9 indicated the copies of RNA per reaction from 4.75 X 101 to 4.74 X 109.

## Discussion

The pandemic of COVID-19 is still going on, particularly with rapid spreading of Omicron and JN.1 (Espinosa-Gongora et al., 2023; Wang, Lu, & Jiang, 2024; Yang et al., 2024). It is expected that new variants of SARS-CoV-2 will have higher transmissibility based on large number of the infected population who may not have symptoms but shed the virus(Kaku et al., 2024; Wannigama et al., 2024). Though some countries stopped enforced COVID-19 test, some hospitals and individuals still need COVID-19 test to help disease prevention and treatment.

In this study, we developed a saliva based, colorimetric RT-LAMP assay that utilized primers targeting the highly conserved SARS-CoV-2 N gene that has been used in other LAMP assays. The detection limit of the developed RT-LAMP assay was 95 copies/test which is close to the sensitivity of the qRT-PCR. Lamb et al. reported their RT-LAMP assay with a sensitivity of 0.2 femtograms but requires ultraviolet light to interpret the results in contrast to our colorimetric assay(Lamb, Bartolone, Ward, & Chancellor, 2020).

Our RT-LAMP assay was comparable to that of qRT-PCR results in terms of sensitivity and specificity. Similar results were described by other labs that reported similar RT-LAMP assays for the detection of SARS-CoV-2[2,4,6]. We employed the direct use of inactivated saliva in the assay without nucleic acid extraction and found out that both qRT-PCR and RT-LAMP produced expected results. Saliva has been demonstrated to be a rich resource for nucleic acids released by microvesicles or viral particles(Balamane et al., 2010; Nagaraj et al., 2006; Robinson, Lee, Kothapalli, Craig, & Fox, 2008). However, due to the viscosity of the saliva and some inhibitors of PCR presented in the saliva, nucleic acids were usually purified from the saliva to not only enrich the nucleic acids, but also eliminate potential inhibitors that might affect downstream processes(Bustos-Garcia et al., 2022; Nishibata et al., 2021). Some reports used enzymes to digest saliva to reduce viscosity and used it directly for nucleic acids analysis(Kim et al., 2024; Rokni et al., 2023). These methods increased the cost and the time of the assay. We tested direct inactivation of the saliva at 65°C for 30 min or 95°C for 5 min and then diluted the saliva with sterile water or PBS. Our data showed that under these conditions, the nucleic acids can be detected easily in RT-PCR or RT-LAMP, reducing the cost and time of the assay. Though the sensitivity of out assay was about 5 to 10-fold less than other methods, it was enough for family use as the sensitivity was enough to screening out the infected individuals based on reported viral RNA in positive saliva(Lalli et al., 2021; Yamazaki, Matsumura, Thongchankaew-Seo, Yamazaki, & Nagao, 2021). The test results can be obtained within 30 minutes from the start of the reaction and its interpretation does not require any sophisticated equipment as it can be easily detected by a change in color from red to yellow.

A major drawback in the current study was that no clinical samples were tested. This was due to the lack of P3 safety lab on the campus. We are expected to test our assay using clinical samples in the future to optimize the protocol.

Based on our data, the colorimetric RT-LAMP would be more suitable as the primary test of family that can identify infected individual with a moderate to high viral load. Moreover, the cost of the assay was calculated to be less than $1.0 when using the master mix at 10 μL compared to $2.3 and over 4 hours. The method described here has many advantages over PCR testing as more people can be screened at a lower cost at home and the results can be obtained much more quickly, enabling earlier identification of infected individuals to help prevent outbreaks of respiratory infectious disease including SARS-CoV-2.

## Conclusions

The developed colorimetric RT-LAMP proved to be highly specific and highly sensitive when compared to qRT-PCR results. Despite the inability to amplify the RNA from clinical samples with a very low viral load at its current state, the test is still suitable to be used as a screening test that can identify the majority of patients with COVID-19. The test could be implemented at workplaces, mobile labs, small clinics, and airports since there is no need for sophisticated equipment and the results can be interpreted immediately with the eye without the need for further analysis. The test proved to be quick and reproducible with robust results as a screening test with reduced cost.

## Acknowledgement

This study was supported by grant 2020SK3034, Dept. of Sci & Tech, Hunan Province, China. The authors declare no conflict of interest.

